# Systematic comparison of developmental GRNs explains how novelty is incorporated in early development

**DOI:** 10.1101/2020.08.07.231464

**Authors:** Gregory A. Cary, Brenna S. McCauley, Olga Zueva, Joseph Pattinato, William Longabaugh, Veronica F. Hinman

**Affiliations:** Department of Biological Sciences, Carnegie Mellon University, Pittsburgh. PA. 15213 USA; The Jackson Laboratory, Bar Harbor, ME, USA; Huffington Center on Aging, Baylor College of Medicine, 1 Baylor Plaza, Houston, TX 77030, USA; Institute for Systems Biology, Seattle, WA, USA

## Abstract

The impressive array of morphological diversity among animal taxa represents the product of millions of years of evolution. Morphology is the output of development, therefore phenotypic evolution arises from changes to the topology of the gene regulatory networks (GRNs) that control the highly coordinated process of embryogenesis^1^. While genetic variation can arise anywhere in the genome and affect any part of an individual GRN, the need to form a viable embryo provides a constraint on the types of variation that pass the filter of selection. A particular challenge in understanding the origins of animal diversity lies in determining how GRNs incorporate novelty while preserving the overall stability of the network, and hence, embryonic viability. Here we assemble a comprehensive GRN, consisting of 42 genes (nodes) and 84 interactions (edges), for the model of endomesoderm specification in the sea star from zygote through gastrulation that corresponds to the GRN for sea urchin development of equivalent territories and stages^2^. Using these detailed models we make the first systems-level comparison of early development and examine how novelty is incorporated into GRNs. We show how the GRN is resilient to the introduction of a transcription factor, *pmar1*, the inclusion of which leads to a switch between two stable modes of Delta-Notch signaling. Signaling pathways can function in multiple modes and we propose that GRN changes that lead to switches between modes may be a common evolutionary mechanism for changes in embryogenesis. Our data additionally proposes a model in which evolutionarily conserved network motifs, or kernels, may function throughout development to stabilize these signaling transitions.

---

The regulatory program that controls development is unidirectional and hierarchical. It initiates with early asymmetries that activate highly coordinated cascades of gene regulatory interactions known as a gene regulatory network (GRN). GRNs function to orchestrate the intricate cellular and morphogenic events that comprise embryogenesis^1,3^, and their topologies must be structured in ways that permit the robust development needed to reliably produce viable embryos. The timing and mechanisms of potential developmental constraint persist as topics of intense debate^4–8^, which can only be resolved by systems-level comparisons of experimentally established GRNs. The evolution of transcription factors (TFs) used in early development presents an especially intriguing problem in the context of maintaining developmental stability^9^.

The GRN for the specification of sea urchin endomesoderm is the most comprehensive, experimentally derived GRN known to date^2,10-11^. It explains how vegetal-most micromeres express signaling molecules, including Delta, needed to specify the adjacent macromere cells to endomesoderm, how micromeres ingress as mesenchyme, and are finally specified to form a biomineralized skeleton. This GRN initiates with the maternally directed nuclearization of β-catenin which activates the paired homeodomain TF *pmar1*^*12*^. Pmar1 represses the expression of *hesC*. The HesC TF is a repressor of genes encoding many of the TFs needed to specify micromere fate (i.e. *alx1, ets1, tbr*, and *tel*) including the *delta* gene. The activation of Pmar1 therefore indirectly leads to the expression of many of the regulatory genes within the vegetal pole, micromere territory in what has been termed the double-negative gate^13^. The Pmar1 TF appears to be a novel duplication of the *phb* gene, and is found only in sea urchins^14^, and the Pmar1 repression of *hesc*, i.e. the double-negative gate^13^, functions only in modern sea urchins. No clear ortholog of *pmar1* exists in available genomes or transcriptomes from sea stars (*Patiria miniata* and *Acanthaster planci*), brittle stars (*Amphiura filiformis*), or hemichordates (*Saccoglossus kowalevskii*)^15–19^ and thus the double-negative gate functions only in sea urchins.

Understanding the impact of integrating this novel subcircuit into early development demands a systems-level approach: not one limited to local properties around the new circuit, but an understanding of how the network as a whole responds to the change. Therefore, we assembled a detailed GRN for sea star endomesoderm specification through gastrulation (Extended Data Figure 1). An interactive, temporal model, including primary and published data, is hosted on a web server (grns.biotapestry.org/PmEndomes) which allows for further and more fine-grained exploration. This GRN was produced using the same experimental approaches as those used to generate the sea urchin network^20^ to allow for a meaningful comparison; i.e., whole-mount in situ hybridization (WMISH) to determine spatiotemporal gene expression, and QRT-PCR and WMISH in control and morpholino antisense oligonucleotide and small molecule inhibitor-induced knockdown of protein function. This network, comprising 42 nodes and 84 edges, approaches in scope the GRN for endomesoderm specification of equivalent stages in sea urchin^2^, at present comprising 72 nodes and 271 edges. This new sea star GRN therefore presents an unprecedented opportunity to compare these networks to understand how they have evolved.

In contrast to the sea urchin^13^ (Figure 1F), sea star blastula stage embryos co-express the *delta* and *hesC* transcripts throughout the endomesoderm-fated vegetal pole (Figure 1A), but their expression is partitioned into adjacent cells by midway through gastrulation (Figure 1B). Indeed, in all known species of echinoderms that lack an identifiable *pmar1* gene, *hesC* remains expressed within the *delta*^*+*^ territory of the blastula mesoderm^15,17,21^. The related Phb genes have recently been shown to act as positive inputs into endomesodermally expressed *hesc* in sea star and cidaroid sea urchin^14^. We show that Tgif also positively regulates *hesC* expression specifically in the sea star mesoderm (Figure 1E). This input is not possible in the sea urchin given that these transcription factors are not co-expressed^22,23^. Not all inputs into *hesC* are changed, however, as *blimp1* (formerly *krox*), a known repressor of *hesC* in sea urchin^24^, also represses sea star *hesC* (Figure 1C-D). Thus the gain of repression by Pmar1 and the loss of positive input from Tgif drive the differences in *hesC* and *delta* co-expression between sea urchins and sea stars (Figure 1G). The local impact of integrating *pmar1* in the sea urchin network, therefore, is the exclusion of *hesC* expression from the territory of cells that will express the *delta* ligand at the vegetal pole. It is via the Delta signal that these cells induce adjacent cells to various endomesodermal fates^25–28^. The early asymmetry in expression of the Delta ligand makes sea urchin micromeres sufficient to induce ectopic endomesoderm when transplanted to the animal pole or to animal blastomeres alone^29^. Thus Delta induction is critical to specifying mesodermal cell types and is one of the central genes providing the regulatory function of the micromeres in sea urchins.

**Figure 1.**
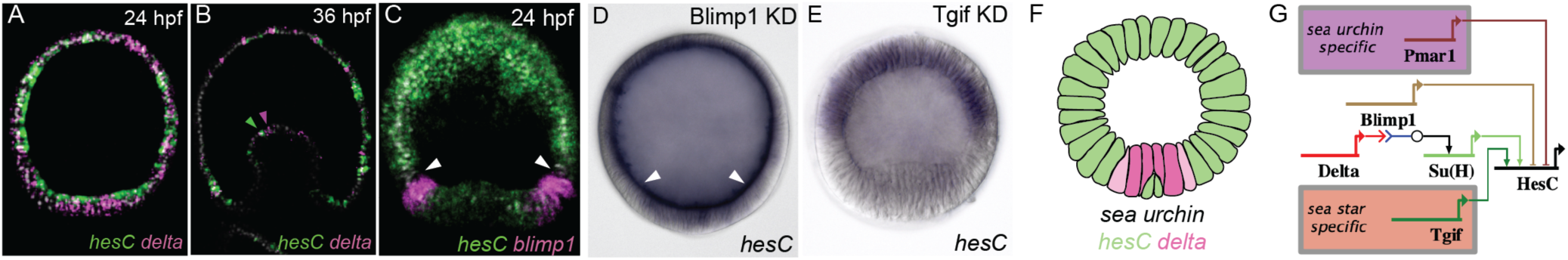
Sea star *hesC* is positively regulated downstream of the mesoderm kernel and is co-expressed with *delta*. Sea star *hesC* and *delta* transcripts are co-expressed in the vegetal mesoderm (A) until mid-gastrula stage (B). Sea star *hesC* and *blimp1* are expressed in partially non-overlapping domains during blastula stage (C) and morpholino knockdown of *blimp1* results in an expansion of the expression domain of *hesC* (D). Morpholino knockdown of sea star Tgif produces a mesoderm-specific decrease in *hesC* expression (E). Schematic showing non-overlapping expression domains of sea urchin *hesC* and *delta* transcripts at blastula stage (F). Regulatory inputs to the *hesC* gene that are specific to sea urchin embryos, sea star embryos, and those that are common to both (G). Data shown are double fluorescent WMISH showing both hesC expression (green) and either expression of delta or blimp1 (magenta) (A-C) and colorimetric WMISH (D-E).

The co-expression of transcripts encoding Delta, HesC, and the Notch receptor (Extended Data Figure 2) in the sea star vegetal pole suggests the potential for lateral inhibition regulatory interactions. In many contexts lateral inhibition functions to segregate an equipotential field of cells into distinct cell types^30–33^. Perturbing Delta-Notch signaling and the expression of *hesC* in sea star blastulae reveals that signaling between adjacent cells through the Notch receptor activates the expression of *hesC*, which in turn represses *delta* expression (Extended Data Figure 3). Thus, we demonstrate that the change in upstream regulation between sea urchin and sea stars that results in co-expression of *delta* and *hesC* at blastula stage allows for lateral inhibition (LI) regulatory interactions in sea stars, compared to the inductive mechanism used in sea urchins (Extended Data Figure 3).

Given that Delta, Notch, and HesC engage in canonical LI, we sought to understand how this process might function in specifying cell types in the sea star mesoderm. The sea star mesoderm originates from the central vegetal pole of the late blastula, a molecularly uniform territory (Figure 2A-B and Extended Data Figure 4). During gastrulation this territory sits at the top of the archenteron and segregates into at least two distinct cell types — blastocoelar mesenchyme and coelomic epithelium (Extended Data Figure 4). These two lineages become molecularly distinct by mid-gastrula stage (approximately 36 hours) and *ets1* expressing mesenchyme cells begin ingressing at 46 hours (Extended Data Figure 5). Each *ets1*^*+*^ cell is generally separated from another *ets1*^*+*^ cell by two intervening nuclei of *ets1*^*-*^ cells while the intervening cells express *six3*, a gene that is also initially broadly mesodermal in blastulae, but is later expressed in the coelomic epithelium (Figure 2C-D and Extended Data Figure 4). *ets1*^*+*^ cells also express the transcript encoding the Delta ligand (Figure 2E). From these data we propose a model in which lateral inhibition leads to the restricted expression of *ets1* in the *delta*^*+*^ cell and *six3* in the neighboring cell.

**Figure 2.**
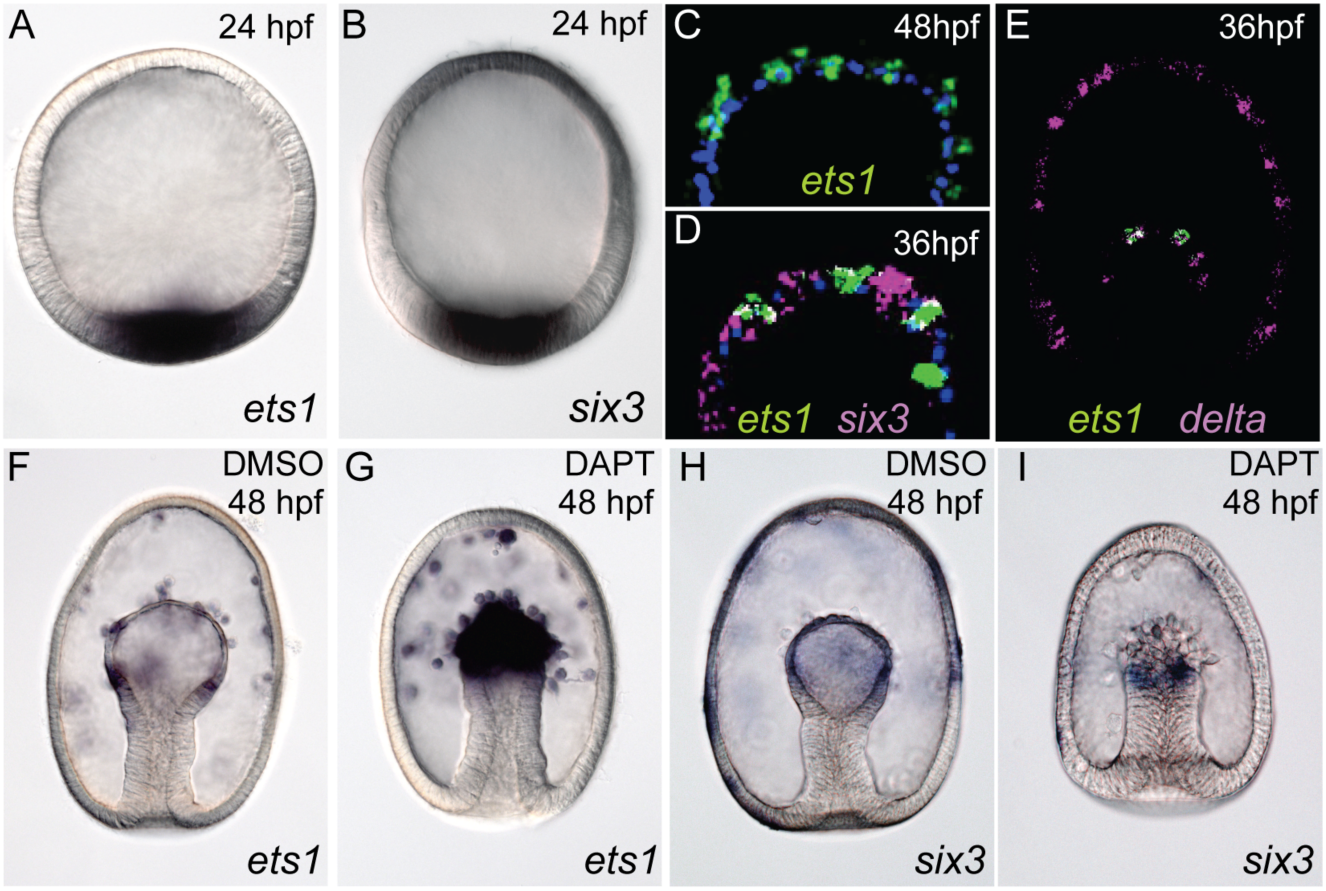
Mesodermal sub-types are specified by Delta-Notch lateral inhibition during gastrulation. Sea star *ets1* and *six3* transcripts are co-expressed in the vegetal mesoderm of blastula stage embryos (A, B). By gastrula stage, cells expressing *ets1* transcript are adjacent to cells with no detectable *ets1* expression (C) and are interleaved by cells expressing *six3* transcript (D). *ets1* expressing cells also express the *delta* transcript (E). Ingressing mesenchyme cells express *ets1* (F) and cells that do not ingress express *six3* (H). Inhibition of Notch with DAPT results in an increase in cells expressing *ets1* (G) and a reduction in cells expressing *six3* (I). There is also a consistent morphological shift with an increase in the number of ingressing cells and a reduction in the epithelium when Notch signalling is blocked. Data shown are colorimetric WMISH (A-B and F-I) and fluorescent WMISH (C-E) with probes indicated in matching colors. Fluorescent images (C-D) are also shown with a nuclear counterstain (DAPI, blue).

We inhibited Notch signaling to explicitly test the lateral inhibition model. The model predicts that such a treatment would lead to an expansion of the primary cell type (*i.e., delta*^*+*^ presumptive mesenchyme) and a concomitant reduction of the secondary cell type (*i.e.*, presumptive coelomic mesoderm). Indeed inhibition of Notch results in an increase in cells expressing the mesenchyme genes *ets1* and *erg*, and a reduction in cells expressing coelomic epithelium genes *six3* and *pax6* (Figure 2F-I and Extended Data Figure 6). Moreover, we also observe a consistent morphological shift with an increase in the number of cells ingressing into the blastocoel from the archenteron and a reduction in the epithelium. These data confirm our hypothesis that the sea star mesoderm partitions through the action of Delta-Notch mediated lateral inhibition. This also shows a consequence of the changes to the regulation of *hesC* that allow coexpression with *delta* — a switch in the mode of Delta-Notch signaling between lateral inhibition when coexpressed and induction when spatially distinct. Asymmetric expression of Delta-Notch regulators typically produces an inductive mode of signaling^34^, and these results suggest that the incorporation of Pmar1 into the early sea urchin network has contributed to this switch.

In sea star embryos, Delta-Notch LI segregates mesenchyme from coelom. While *hesC* expression is associated with cells fated to the coelom, *hesC* expression is no longer detected in the mesoderm by 48 hours (Extended Data, Figure 7), shortly after the completion of cell type partitioning and coincident with the onset of epithelial-mesenchymal transition. The Delta-Notch LI must then “hand-off” to another set of genes to stabilize and maintain coelomic restricted gene expression. Correspondingly, this stage is also the first time that we observe the expression of *pax6* and *eya* in this territory^35^. *pax6, eya, six1/2, dach*, and *six3* are expressed in coelomic-fated mesoderm in both sea stars and sea urchins along with other genes from the highly conserved retinal determination gene network (RDGN)^35–36^. In the sea urchin coelom, these genes are wired together in a network topology considered to be homologous to that of the RDGN ^37^. We examined the regulatory interactions between *pax6, six3, eya, dach*, and *six1/2* to determine if a similar network is involved in the maintenance of sea star coelomic mesoderm. We find a similar activation of *six3* by Pax6, of *six1/2* by itself, and of *eya* by Pax6, Six3, and Dach (Extended Data Figure 8). Thus, these genes interact in a similar regulatory sub-network in both sea star and sea urchin coelomic mesoderm, and this sub-circuit is highly similar to the *Drosophila* RDGN (Extended Data Figure 8), suggesting even deeper conservation of this network architecture. While Delta-Notch mediated lateral inhibition is responsible for early segregation of mesenchymal and coelomic cell fates in sea stars, a Pax6 and Six3 mediated network is necessary for proper coelomogenesis after the completion of lateral inhibition (Extended Data Figure 7).

The summary view of these results permits a global comparison of these echinoderm GRN topologies (Figure 3A), which are the synthesis of over a decade of work including the present study. Immediately apparent are the several distinct subcircuits found in common between these networks. Common modules include *ets1, erg, hex*, and *tgif* in the mesoderm^38,39^; *otx, gatae, bra*, and *foxa* in the endoderm^40,41^; and *pax6, eya, six3, dach*, and *six1/2* in the coelom (Extended Data Figure 8, ^37^). In contrast, entire subcircuits present in sea urchins, *i.e. dri, foxb*, and *vefgr* that direct batteries of skeletogenic differentiation genes in the sea urchin micromeres, are entirely absent from the sea star network. This comparison also reveals that similarly regulated subcircuits are highly positively cross regulated, in keeping with the previous definition of network kernel^42^. It was previously suggested that such kernels would be found in early development, as they function downstream of maternally derived and transient signals, to stabilize gene expression needed to specify distinct embryonic territories^43^. A stabilizing function is thought to be derived from the intra-circuit positive regulatory feedback. Here we show that these kernels appear throughout the GRN and are not limited to only early development. Preliminary experimental analyses from other species of echinoderms^15,44–46^ suggest these kernels are present in multiple species and thus represent a genuinely conserved, rather than convergent, feature of GRNs. The mechanistic basis for the evolutionary stability of these subcircuits remains unclear and it will be important to define additional such network motifs to begin to understand whether the observed stability is a cause or a consequence of the observed highly recursive regulatory wiring of these motifs^42^.

**Figure 3.**
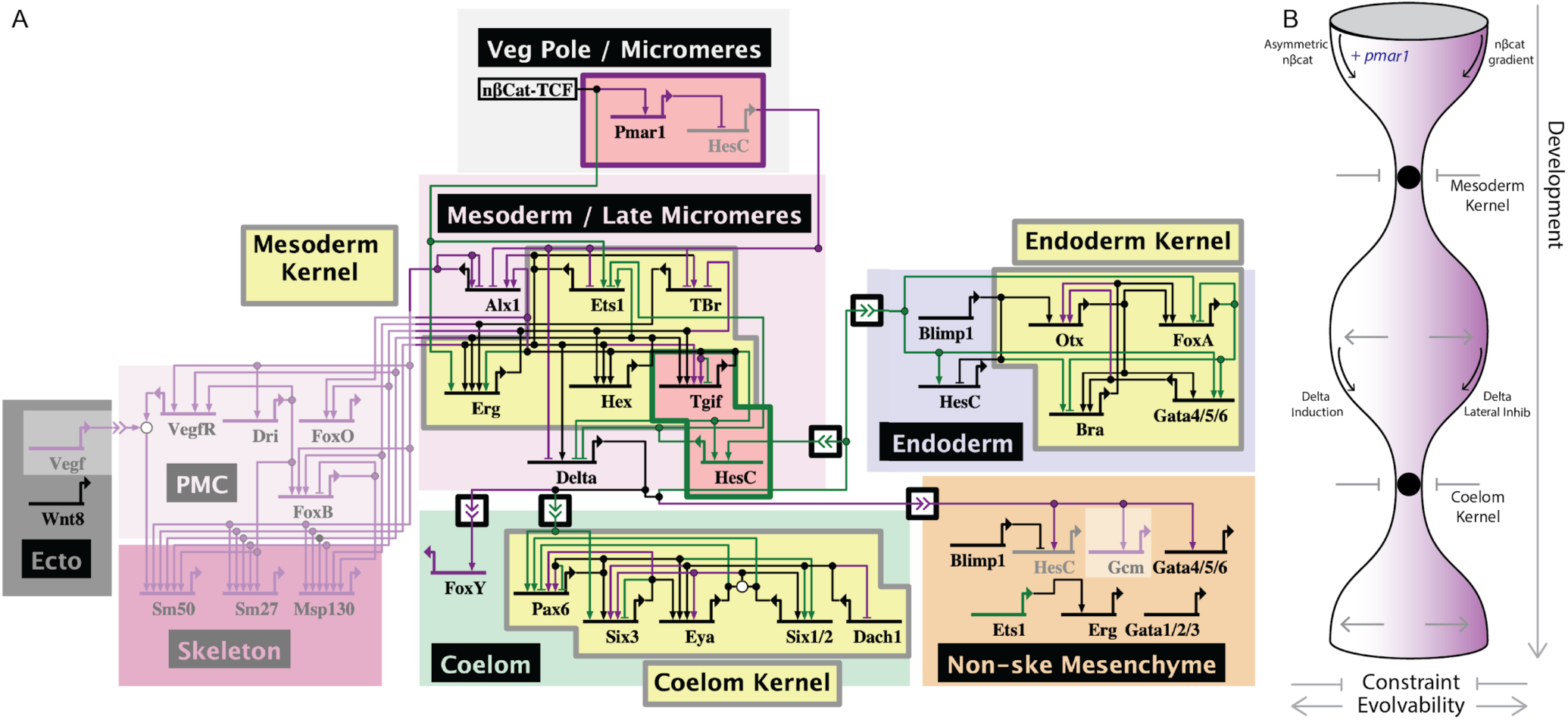
Evolutionary constraint of network kernels permits alterations to the surrounding network. A synthesis of key aspects of the GRN from sea urchins and sea stars is shown (A). Genes (nodes) are shown in the territories (colored boxes) in which they are expressed. Edges are regulated by the originating upstream factor and are either positive (arrow) or repressive (bar). Signaling across cell types are indicated as double arrow heads, and Delta to Notch signals are boxed. Genes and links that are unique to sea urchin embryos are colored purple, those specific to sea stars are green, and those in common are black. Network kernels are highlighted (yellow) as are distinct sub-circuits (pink), including the sea urchin specific double negative gate (i.e. Pmar and HesC, purple outline) and sea star specific positive regulation of HesC (i.e. Tgif and HesC, green outline). Greyed out backgrounds indicate entire network circuits that are absent in sea stars. Our model of GRN evolution is depicted (B) showing that network kernels are constrained regions whereas both up and down the hierarchy the network is capable of change.

From these data we propose a model of how changes in the GRN are incorporated while maintaining an overall network stability. We have detailed how the network incorporates novel circuitry into early development, in this example, the Pmar1*-*HesC double-negative gate. These networks use the same signaling pathways at the same places in the GRN, but rely on different signaling modalities of the pathways; the networks use binary versus dosage dependence of nβ-catenin^47^, and Delta induction versus Delta-Notch lateral inhibition. We argue that changes in GRNs, such as the introduction of novel genes or subcircuits, that lead to switching between alternate, stable modes of signaling pathways may be a common source of evolutionary change in these GRNs. The disruption caused by such changes would be limited if they are surrounded by stabilization features. Indeed, despite this transition in Delta-Notch signaling, we find that the GRNs in both taxa converge to a conserved regulatory subcircuit that directs the fate of coelomic mesoderm. Therefore, in contrast to previous expectations, evolutionarily stable network kernels that were proposed to function to lockdown early developmental regulatory states^3^, are not restricted according to network hierarchy. We propose instead that these network subcircuits act as stabilizing features throughout development, functioning as developmental checkpoints through which embryogenesis must transit (Figure 3B).

## Supporting information

BioTapestry btp file

