## Supplemental methods, figures, and tables S1-S3 for "Systematic comparison of developmental GRNs explains how novelty is incorporated in early development"

*Animal and embryo handling*

Adult *Patiria miniata* were obtained from the southern coast of California, USA (Pete Halmay or Marinus Scientific) and were used to initiate embryo cultures as previously described[1]. *P. miniata* embryos were cultured in artificial seawater at 16 °C.

*Whole-Mount Staining*

Embryos were fixed as previously described[2,3]. Briefly, embryos were fixed in a solution of 4% paraformaldehyde in MOPS-fix buffer (0.1M MOPS pH 7.5, 2mM $MgSO_4$, 1mM EGTA, and 800 mM NaCl) for 90 minutes at 25 °C and transferred to a solution of 70% ethanol for long term storage at -20 °C. In situ hybridization experiments were performed as previously described[3,4] using digoxigenin-labeled antisense RNA probes. Probes were designed using gene model sequence predictions from Echinobase.org [5,6]. For two-color FISH a second dinitrophenyl-labeled antisense RNA probe was hybridized simultaneously[3]. Images of colorimetric whole mount specimens were taken using a Leica DMI 4000B microscope equipped with a Leica DFC 420C camera and fluorescent specimens were photographed using a Zeiss LSM 880 scanning laser confocal microscope. At least two independent biological replicate experiments were performed for each in situ staining experiment, examining the pattern of at least 10 specimens per replicate.

*Quantitative PCR*

RNA was extracted using the GenElute Mammalian Total RNA Kit (Sigma-Aldrich) and DNA was removed using the DNA-free™ DNA Removal Kit (Invitrogen). Quantitative real-time PCR was performed as previously described using the qScript One-Step SYBR Green qRT-PCR Kit (QuantaBio) and the and run on an Applied Biosystems 7300 Real-Time PCR instrument. Measured Ct values for reported genes were normalized to the Ct of an internal control *lamin2β receptor* (GenBank ID: KJ814251.1;[7]). The sequence of all qPCR primers used is reported in Extended Data Table S2.

*Perturbation of gene expression*

Zygotes were injected with morpholino antisense oligonucleotides (MASOs; GeneTools) following the protocol by Cheatle Jarvela and Hinman[1]. For all MASOs, the GeneTools standard control MASO was injected into sibling embryos. The observed phenotype of each MASO knockdown was confirmed by injecting a second MASO designed to the same transcript. The sequence and effective concentration used for each MASO used is reported in Extended Data Table S1. Notch perturbations were achieved by bathing embryos in 32 μM DAPT[8] or DMSO as a control from the two-cell stage. WMISH was performed on at least three independent sets of perturbed embryos. At least ten embryos were assessed in each replicate and phenotypes were counted and a summary is reported in Extended Data Table S3. Quantitative measures of perturbation were achieved by performing qPCR on perturbed compared with control siblings. Each assay was performed on at least two qPCR replicates in each of two biological replicates.

*Gene Regulatory Network Construction*

The GRN model depicting sea star endomesoderm was constructed using BioTapestry[9,10]. The network was constructed by reviewing literature, spanning the years 2003-2019, which describes both embryonic gene expression and gene regulation in *Patiria miniata*. The experimental evidence supporting each node and edge is provided in Extended Data Table S4 and all references to work cited in the GRN experimental evidence utilized are herein cited[2,3,7,11–23]. The expression and regulatory linkages are included as reported and have been ordered according to the embryonic chronology and spatially arranged into appropriate territories. A summary of these findings is presented in Extended Data Figure 1, and a dynamic and interactive model of a more detailed network is hosted on the web at http://grns.BioTapestry.org/PmEndomes/ for further and more fine-grained exploration of the GRN[24]. Additionally, a BioTapestry .btp file is included in the supplemental materials and can be viewed using the BioTapestry desktop application available for download at http://www.BioTapestry.org. We expect that the online model will continue to be updated in the future to capture changes to the network.

There are three principal temporal subdivisions that span 10-24 hpf, 25-34 hpf, and 35-50 hpf, the breakpoints relating to major embryonic milestones, i.e. the distinction of mesoderm from endoderm at ~24 hpf and subsequently the split between mesenchymal and coelomic fated mesoderm at ~35 hpf. The BioTapestry user can select which of these models to view by clicking on it in the left-hand panel of the viewer. The models are organized in a hierarchy, with the top-level Full Genome model showing all nodes and links present in all the submodels. The three submodels, representing the three principal temporal subdivisions listed above, summarize the behavior of the network in the various modeled developmental domains that exist for that time period. Below each of these models in the hierarchy are dynamic models that show hourly views (using the time slider in the lower left) for that period. Note that though the time slider is hourly, expression states between the experimental data points (five hours apart) are being interpolated.

The BioTapestry model is designed to summarize the known information about the *P. miniata* developmental GRN, obtained both from literature and from our experimental results, and it is crucial to use the interactive online version to best understand the behavior of the network. The differential temporal and spatial expression patterns of the genes in the network, as determined by experiment and known with a high degree of confidence, are depicted by showing the genes as “on” or “off” (colored or grey, respectively) in the various regions of the model at each timepoint. In nodes where variable levels of expression are crucial (viz. the gradient of nuclearized beta catenin/TCF from Mesoderm to Veg1 Ectoderm that is present in the early Endomesoderm 10-24 hour summary model) the nodes are depicted using intermediate levels of grey to colored. Note that by right-clicking on a gene name and selecting Experimental Data from the pop-up menu, you can view the underlying experimental expression data for the gene as well as experimental data supporting inputs to the gene. The edges of the network are also based upon literature and experimental results. Of course, there are many different levels of confidence that can be assigned to each edge, based upon the type of experiment, and colored diamonds below the link terminus on target genes are used to indicate confidence. The highest confidence based upon detailed cis-regulatory analysis of a target gene[21], is depicted with a green diamond, see e.g. Tbr activation of *otx* in the GRN model. However, most links in the network are backed by the results of perturbation experiments (e.g. MASO knockdown of the source gene or drug perturbation of a signaling pathway). To ensure only high confidence links are included, we use a threshold of at least 2-fold change observed in a minimum of 2 independent perturbation experiments. We also utilize multiple MASOs targeting the same transcript to ensure specificity of the observed phenotypes (see Extended Data Table S1). While there is no guarantee that these links are in fact direct, direct edges that can be explained through an indirect path can be omitted through a parsimonious approach to adding links to the network.

Links, like nodes, are also shown as “on” or “off” in the model at each point in space and time, simply based on the expression of the source gene at that same point. Notably, this depiction says nothing explicit about the actual cis-regulatory logic that is encoded in the target gene. Just because a link is shown as colored and incident on a target gene does not mean that it has been shown to be necessary at that point in space and time to cause expression of the target gene. To make that conclusion, much more targeted experiments are required to make that claim. However, the on/off state of the target gene and the inbound links can provide clues to generate hypotheses and suggest further experiments. For example, if all the links into a grey (off) target are colored (on), that suggests that there must be other unknown inputs into the target gene.

This model is not purporting to be complete, but is instead a systems-level summary of the existing state of knowledge about the causal mechanisms underpinning *P. miniata* development driven by the GRN. It is certainly missing genes, and in fact since it is heavily based on orthology to genes present within the sea urchin GRN, we expect this network is biased towards including just those transcription factors. Furthermore, it has not been validated by computational simulations, and involves no detailed modeling of the transcriptional mechanisms that control gene expression. In this regard, it is like the sea urchin network, which was first developed using gene expression and perturbation data [25] many years before boolean simulations were performed to ascertain the ability of the model to explain the observed behavior [26].

**Methods references**


1. Cheatle Jarvela, A. M. & Hinman, V. A method for microinjection of Patiria miniata zygotes. *J. Vis. Exp.* e51913 (2014). doi:10.3791/51913
2. McCauley, B. S., Weideman, E. P. & Hinman, V. F. A conserved gene regulatory network subcircuit drives different developmental fates in the vegetal pole of highly divergent echinoderm embryos. *Dev. Biol.* **340,** 200–208 (2010).
3. Yankura, K. A., Martik, M. L., Jennings, C. K. & Hinman, V. F. Uncoupling of complex regulatory patterning during evolution of larval development in echinoderms. *BMC Biol.* **8,** 143 (2010).
4. Hinman, V. F., Nguyen, A. T. & Davidson, E. H. Expression and function of a starfish Otx ortholog, AmOtx: a conserved role for Otx proteins in endoderm development that predates divergence of the eleutherozoa. *Mech. Dev.* **120,** 1165–1176 (2003).
5. Kudtarkar, P. & Cameron, R. A. Echinobase: an expanding resource for echinoderm genomic information. *Database (Oxford)* **2017,** (2017).

6. Cameron, R. A., Samanta, M., Yuan, A., He, D. & Davidson, E. SpBase: the sea urchin genome database and web site. *Nucleic Acids Res.* **37,** D750-4 (2009).
7. McCauley, B. S., Akyar, E., Filliger, L. & Hinman, V. F. Expression of wnt and frizzled genes during early sea star development. *Gene Expr Patterns* **13,** 437–444 (2013).
8. Materna, S. C. & Davidson, E. H. A comprehensive analysis of Delta signaling in pre-gastrular sea urchin embryos. *Dev. Biol.* **364,** 77–87 (2012).
9. Longabaugh, W. J. R., Davidson, E. H. & Bolouri, H. Computational representation of developmental genetic regulatory networks. *Dev. Biol.* **283,** 1–16 (2005).
10. Longabaugh, W. J. R., Davidson, E. H. & Bolouri, H. Visualization, documentation, analysis, and communication of large-scale gene regulatory networks. *Biochim. Biophys. Acta* **1789,** 363–374 (2009).
11. McCauley, B. S., Wright, E. P., Exner, C., Kitazawa, C. & Hinman, V. F. Development of an embryonic skeletogenic mesenchyme lineage in a sea cucumber reveals the trajectory of change for the evolution of novel structures in echinoderms. *Evodevo* **3,** 17 (2012).
12. McCauley, B. S., Akyar, E., Saad, H. R. & Hinman, V. F. Dose-dependent nuclear β-catenin response segregates endomesoderm along the sea star primary axis. *Development* **142,** 207–217 (2015).
13. Gildor, T., Cary, G. A., Lalzar, M., Hinman, V. F. & Ben-Tabou de-Leon, S. Developmental transcriptomes of the sea star, Patiria miniata, illuminate how gene expression changes with evolutionary distance. *Sci. Rep.* **9,** 16201 (2019).
14. Annunziata, R., Martinez, P. & Arnone, M. I. Intact cluster and chordate-like expression of ParaHox genes in a sea star. *BMC Biol.* **11,** 68 (2013).
15. Fresques, T. M. & Wessel, G. M. Nodal induces sequential restriction of germ cell factors during primordial germ cell specification. *Development* **145,** (2018).
16. Fresques, T., Zazueta-Novoa, V., Reich, A. & Wessel, G. M. Selective accumulation of germ-line

associated gene products in early development of the sea star and distinct differences from germ-line development in the sea urchin. *Dev. Dyn.* **243,** 568–587 (2014).

17. Hinman, V. F. & Davidson, E. H. Expression of a gene encoding a Gata transcription factor during embryogenesis of the starfish Asterina miniata. *Gene Expr Patterns* **3,** 419–422 (2003).
18. Hinman, V. F. & Davidson, E. H. Expression of AmKrox, a starfish ortholog of a sea urchin transcription factor essential for endomesodermal specification. *Gene Expr Patterns* **3,** 423–426 (2003).
19. Hinman, V. F. & Davidson, E. H. Evolutionary plasticity of developmental gene regulatory network architecture. *Proc. Natl. Acad. Sci. USA* **104,** 19404–19409 (2007).
20. Hinman, V. F., Nguyen, A. T., Cameron, R. A. & Davidson, E. H. Developmental gene regulatory network architecture across 500 million years of echinoderm evolution. *Proc. Natl. Acad. Sci. USA* **100,** 13356–13361 (2003).
21. Hinman, V. F., Nguyen, A. & Davidson, E. H. Caught in the evolutionary act: precise cis-regulatory basis of difference in the organization of gene networks of sea stars and sea urchins. *Dev. Biol.* **312,** 584–595 (2007).
22. Otim, O., Hinman, V. F. & Davidson, E. H. Expression of AmHNF6, a sea star orthologue of a transcription factor with multiple distinct roles in sea urchin development. *Gene Expr Patterns* **5,** 381–386 (2005).
23. Yankura, K. A., Koechlein, C. S., Cryan, A. F., Cheatle, A. & Hinman, V. F. Gene regulatory network for neurogenesis in a sea star embryo connects broad neural specification and localized patterning. *Proc. Natl. Acad. Sci. USA* **110,** 8591–8596 (2013).
24. Paquette, S. M., Leinonen, K. & Longabaugh, W. J. R. BioTapestry now provides a web application and improved drawing and layout tools. [version 1; peer review: 3 approved]. *F1000Res.* **5,** 39 (2016).
25. Davidson, E. H. *et al.* A genomic regulatory network for development. *Science* **295,** 1669–1678

(2002).

26. Peter, I. S., Faure, E. & Davidson, E. H. Predictive computation of genomic logic processing functions in embryonic development. *Proc. Natl. Acad. Sci. USA* **109,** 16434–16442 (2012).

**Acknowledgements**

The authors thank Dr. William Hatleberg for helpful feedback during the preparation of the manuscript. This work was supported by the Binational Science Foundation grant number 2015031 to VFH, the National Science Foundation grants IOS 1557431 and MCB 1715721 to VFH and the National Institute of Health grant P41HD071837 to VFH.

**Author Contributions**

GC, BM, and VH conceived of and designed experiments. GC, BM, OZ and JP carried out experiments. GC and BM analyzed data and VH was instrumental in the interpretation of the results. GC and WL constructed the network model. GC wrote the manuscript with significant input from BM and VH. All authors read and approved of the final manuscript.

**Competing Interest Declaration**

The authors declare no competing interests.

**Corresponding Author**

Veronica Hinman

(412) 268-9348

### Extended Data - figures, tables, and legends

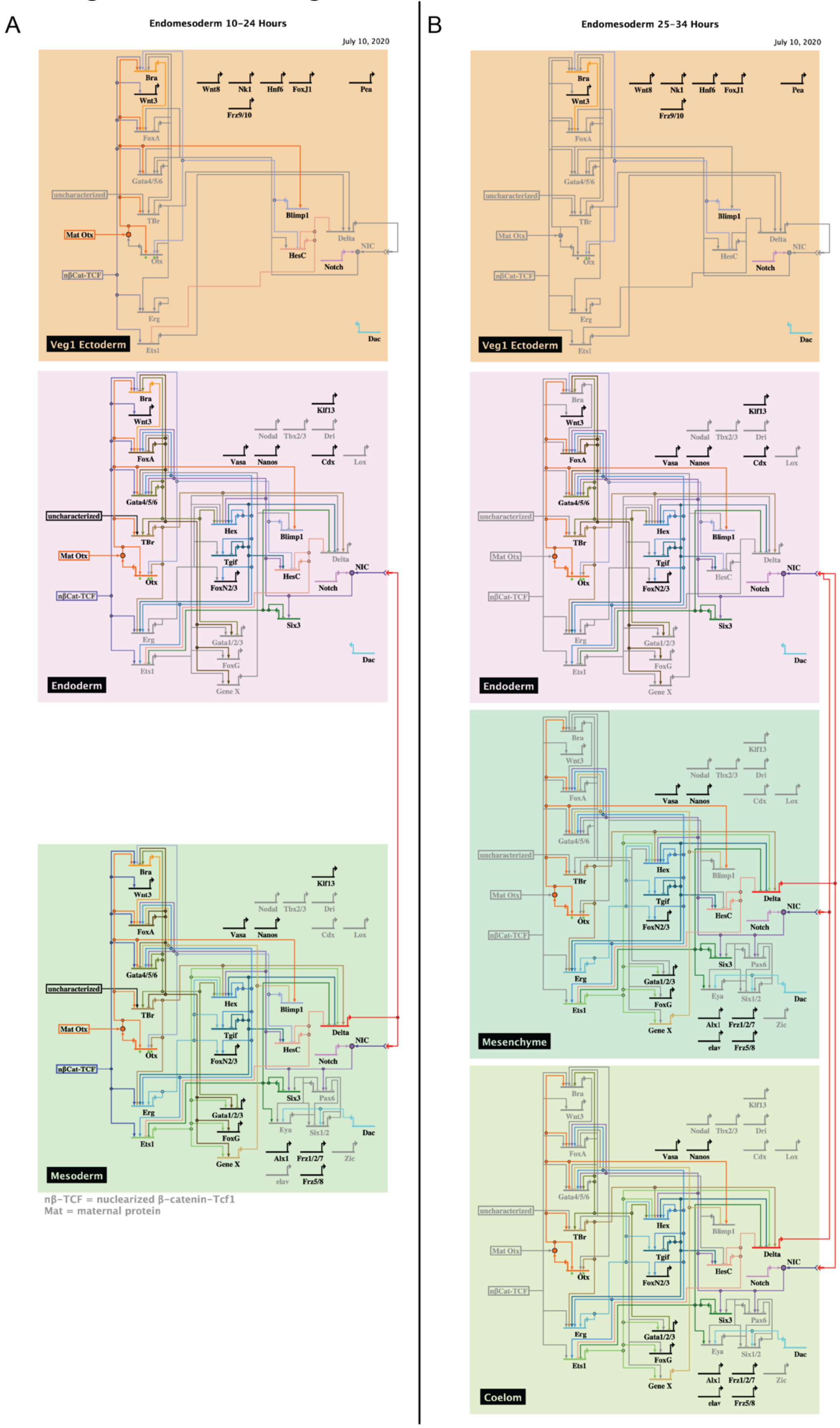


(continued next page)

C

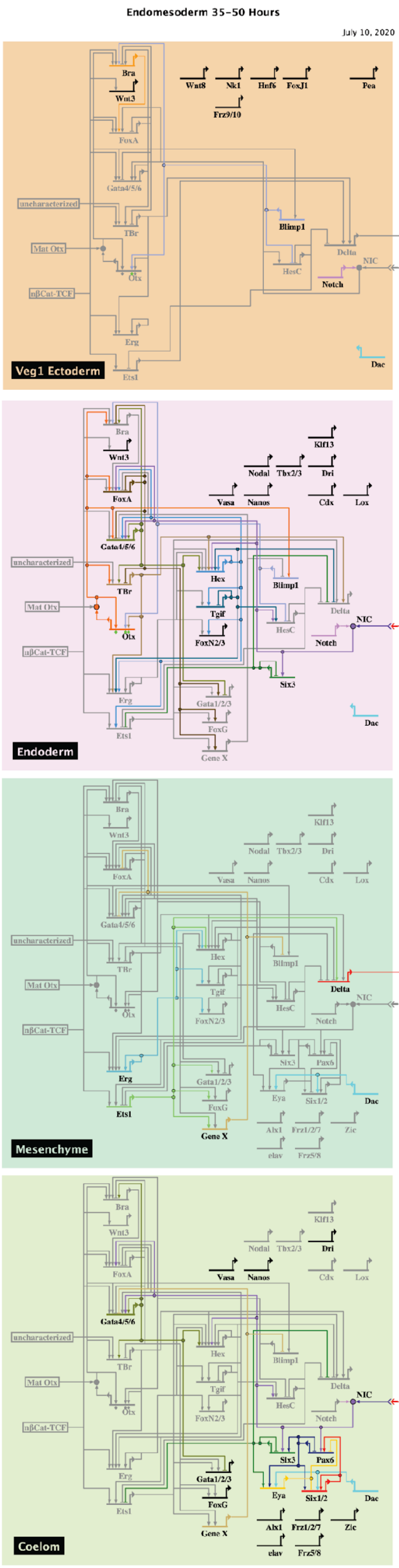
Endomesoderm 35-50 Hours
July 10, 2020
Veg1 Ectoderm
Endoderm
Mesenchyme
Coelom
Bra
Wnt3
FoxA
Gata4/5/6
TBr
Otx
Erg
Ets1
Blimp1
Delta
Notch
NIC
HesC
Hex
Tgif
FoxN2/3
Six3
Gata1/2/3
FoxG
Gene X
Dac
Kif13
Nodal
Tbx2/3
Dri
Vasa
Nanos
Cdx
Lox
Pax6
Eya
Six1/2
Alx1
Frz1/2/7
Zic
elav
Frz5/8
Wnt8
Nk1
Hnf6
FoxJ1
Pea
Frz9/10
uncharacterized
Mat Otx
nβCat-TCF

D

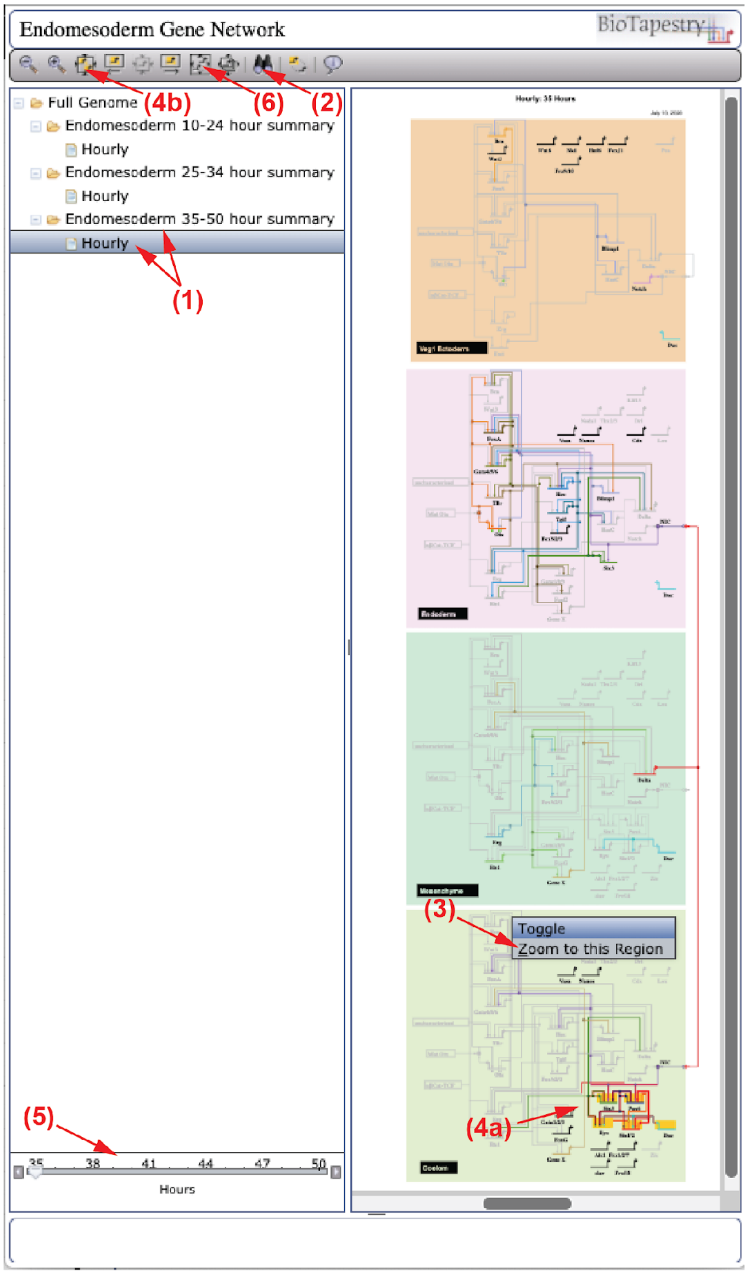
Endomesoderm Gene Network
BioTapestry
Full Genome
Endomesoderm 10-24 hour summary
Hourly
Endomesoderm 25-34 hour summary
Hourly
Endomesoderm 35-50 hour summary
Hourly
(1)
(2)
(3)
(4a)
(4b)
(5)
(6)
Toggle
Zoom to this Region
35
38
41
44
47
50
Hours

**Figure 1.** *P. miniata* endomesodermal network through 50 hours of embryogenesis. These images represent screenshots of the summary views of the extended network available online (grns.biotapestry.org/PmEndomes). The nodes of the network are genes and the horizontal lines represent cis-regulatory input regions into each gene. The nodes are connected by edges that indicate either positive (arrow) or repressive (bar) relationships between two specific genes. Edges with experimental cis-regulatory support are indicated (green diamonds). Genes are grouped into territories (colored boxes) in which they are expressed during specific developmental stages. Signaling between cell types is indicated as double arrow heads. The expression and regulatory linkages are included here as reported and have been ordered according to the embryonic chronology and spatially arranged into appropriate territories. There are three principal temporal subdivisions that span 10-24 hpf (A), 25-34 hpf (B), and 35-50 hpf (C), the breakpoints relating to major embryonic milestones, *i.e.* the distinction of mesoderm from endoderm at ~24 hpf and subsequently the split between mesenchymal and coelomic fated mesoderm at ~35 hpf. An extended, interactive version with dynamic, hour-by-hour views of embryonic progression as well as detailed information about experimental evidence supporting each node and edge assignment is available (grns.biotapestry.org/PmEndomes). Navigation through the online version of the model (D) involves selection of either the summary view or the hourly view (1) of a specific submodel. Genes within the submodel can be identified by searching (2, binocular icon). Higher resolution views of each region can be accessed by zooming to a particular region (3) or highlighting groups of genes (4a) and magnifying the highlighted nodes (4b). The temporal aspect of hourly models can be examined using the time slider (5). Returning to the entire sub-model view can be achieved by clicking on the icon indicated (6). To access information about the experimental data supporting each node or edge, one can click on a network element and right click to bring up a contextual menu from which one can select "Experimental Data". The experimental data supporting the construction of this model is also available as a supplemental table to this manuscript (Extended Data, Table S4).

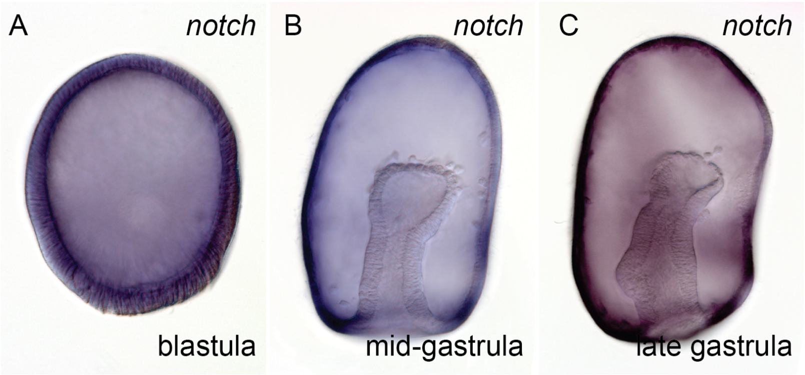


**Figure 2.** Expression of the sea star *notch* transcript by colorimetric WMISH. RNA encoding the Notch receptor is ubiquitously distributed at blastula stage (A) and is restricted to ectodermal expression at mid- and late-gastrula staged (B,C).

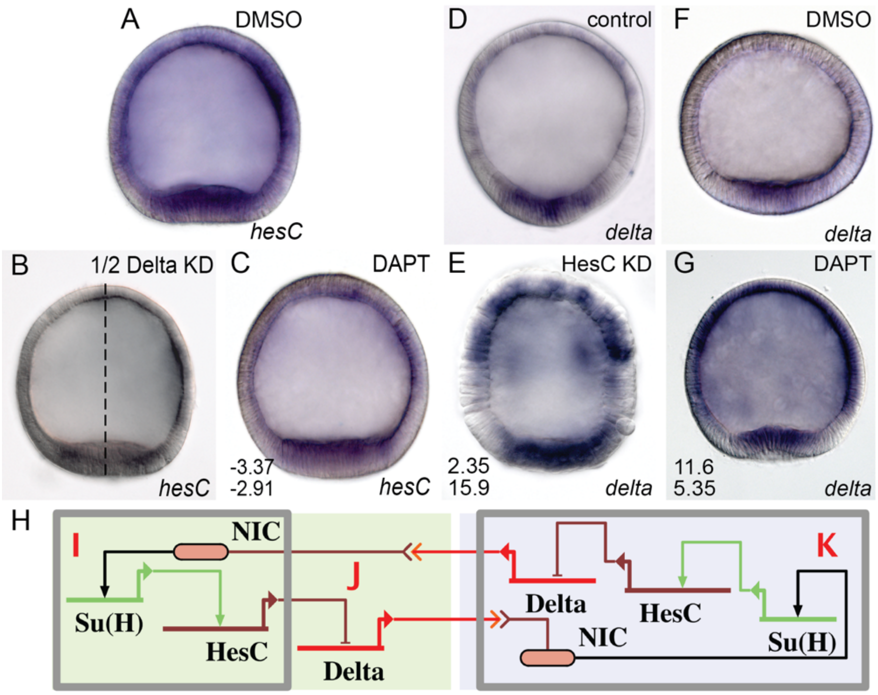


**Figure 3.** Testing lateral inhibition of *delta* and *hesC* by inhibition of Notch signaling (DAPT) and morpholino knockdown of HesC. Using DAPT, an inhibitor of the proteolytic gamma-secretase necessary for notch signal transduction, we observe both a down-regulation of *hesC* (C) and an up-regulation of *delta* transcripts (G). Importantly the down-regulation of *hesC* is phenocopied by injection of a morpholino targeting the *delta* transcript into one of the first two blastomeres (B). Knockdown of HesC with an antisense morpholino yields an up-regulation of both *delta* (E) and *hesC* transcripts. Lateral inhibition network showing relationships tested by previous experiments; red letters indicate figure panel supporting connection (L). All images are colorimetric WMISH with the probes to the indicated genes.

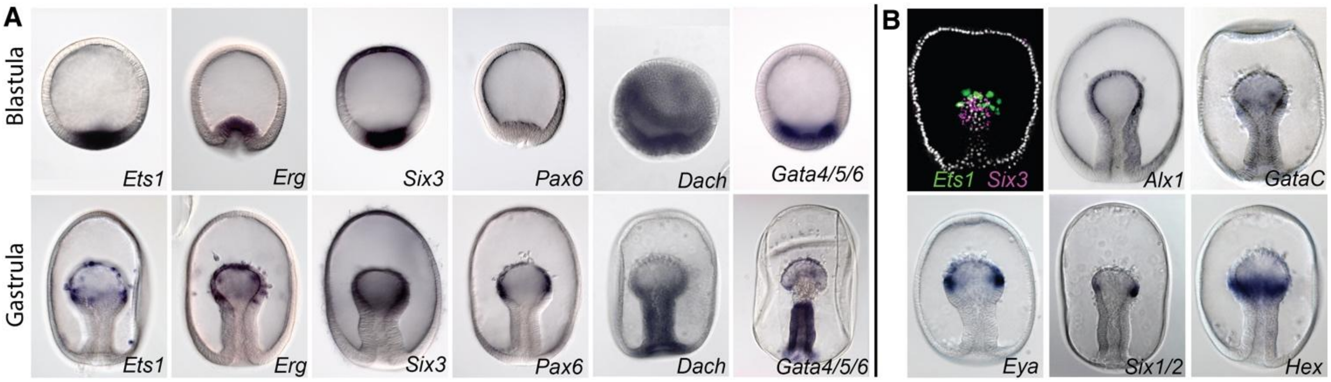


**Figure 4.** Molecularly uniform vegetal pole mesoderm partitions into blastocoelar mesenchyme and coelomic epithelium by mid-gastrula stage by colorimetric WMISH (A). The *ets1*, *erg*, *six3*, *dach,* and *gata4/5/6* transcripts are all expressed in the vegetal pole territory in blastula stage embryos in overlapping domains. The *gata4/5/6* transcript is distributed slightly broader owing to the fact that it is also expressed in the endoderm and the *dach* transcript is expressed ubiquitously early. Expression of *pax6* is not detected until later stages. By mid-gastrula stage both *ets1* and *erg* are expressed in ingressing mesenchyme cells, while *six3*, *pax6*, *dach*, and *gata4/5/6* are all expressed in the mesodermal epithelium at the top of the archenteron. Both *dach* and *gata4/5/6* are also both expressed throughout the endoderm at this stage. Additional ceolomic mesoderm genes are expressed in the posterior aspect of the mesodermal bulb of the archenteron by mid-gastrula stage (B). The expression of *eya*, *six1/2*, *alx1*, and *hex* transcripts are all co-localized in this territory. Importantly, as shown by dFISH, *ets1* expressing cells are distinct from *six3* expressing cells at this stage.

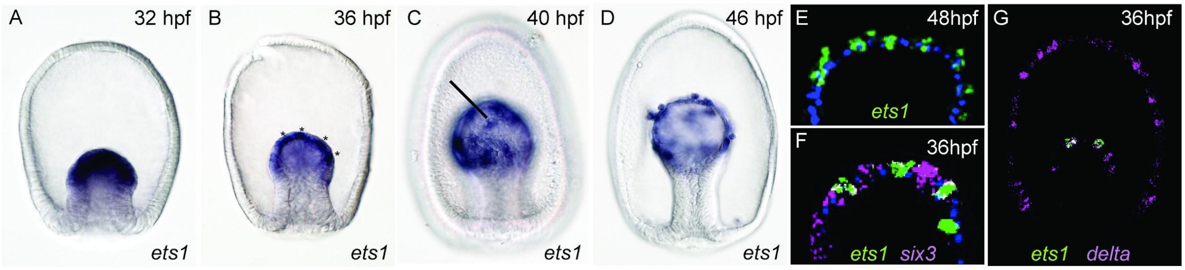


**Figure 5.** Segregation of mesodermal subtypes into interleaved cells by 36 hpf. The expression of the *ets1* transcript was assessed every 2 hours from the onset of gastrulation by colorimetric WMISH. At 32 hpf the expression of *ets1* is uniform throughout the mesoderm (A). At 36 hpf there is a discontinuity in the expression of *ets1* transcript (B, asterisks). Patches of *ets1* expression become more distinct by 40-42 hpf (C), and *ets1* expressing cells start to ingress beginning at 46 hpf (D). Cells expressing *ets1* are adjacent to cells with no detectable *ets1* expression (E), using fluorescent WMISH with a DAPI counterstain (blue). Intervening *ets1*$^{-}$ cells express *six3* (F) by double fluorescent WMISH. Cells expressing *ets1* transcript also express the transcript encoding the Delta ligand (G).

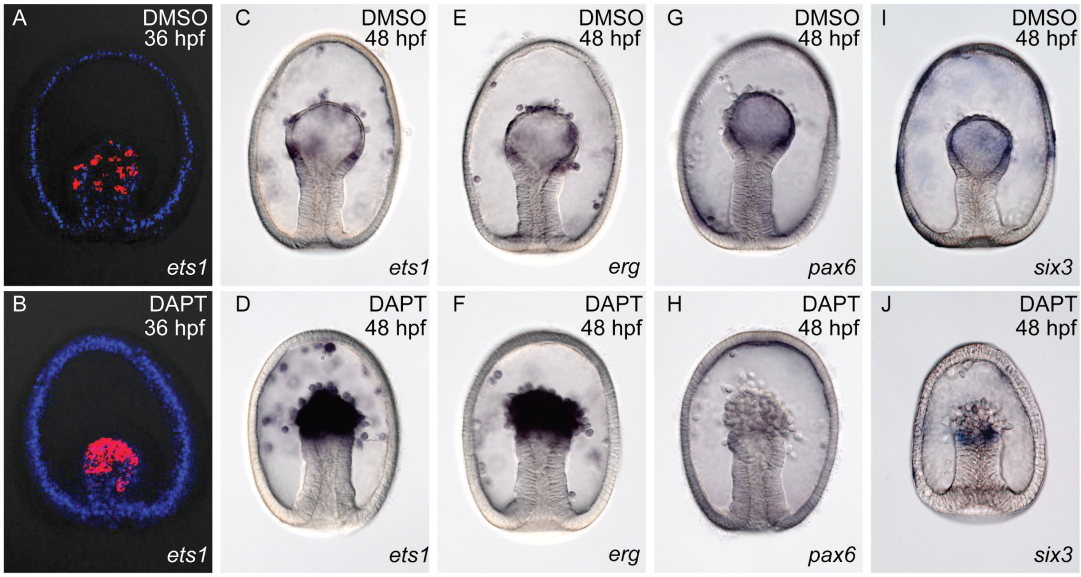


**Figure 6.** Testing the lateral inhibition model of mesodermal subtype segregation. The expression of *ets1* transcript appears in a salt-and-pepper distribution throughout the mesoderm at 36 hpf by fluorescent WMISH (A) with nuclei stained (blue). Treatment with the Notch inhibitor DAPT beginning at the 2-cell stage results in a uniform expression of *ets1* in this territory (B). By 48 hpf, the mesenchyme cells expressing *ets1* and *erg* (C, E) ingress into the blastocoel while cells that do not ingress express *pax6* and *six3* (G, I). DAPT treatment results in an increase in cells expressing *ets1* and *erg* (D, F) and a reduction in cells expressing *pax6* and *six3* (H, J). Data shown in (C-J) are colorimetric WMISH.

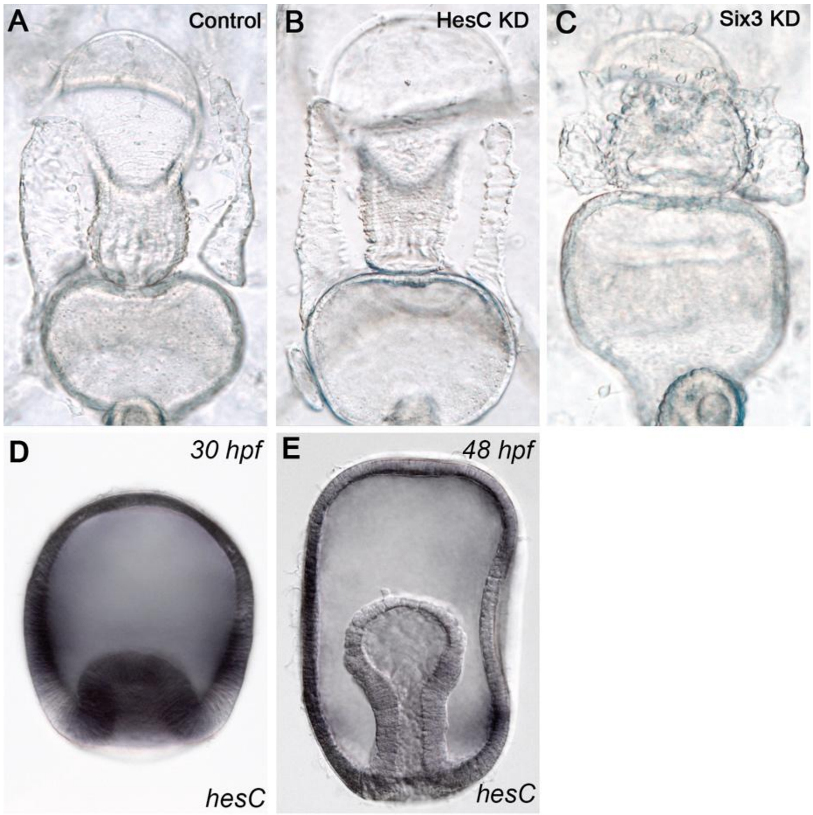


**Figure 7.** HesC handoff by 48 hpf. MASOs targeting *hesC* do not perturb the morphology of the coelomic epithelium at 6 days post fertilization (A, B), while perturbation of *six3* leads to much diminished coelomic epithelium as well as a shortened foregut (C). The expression of *hesC* transcript, detected in early mesodermal territories by colorimetric WMISH (D), is undetectable in mesoderm by 48 hpf (E).

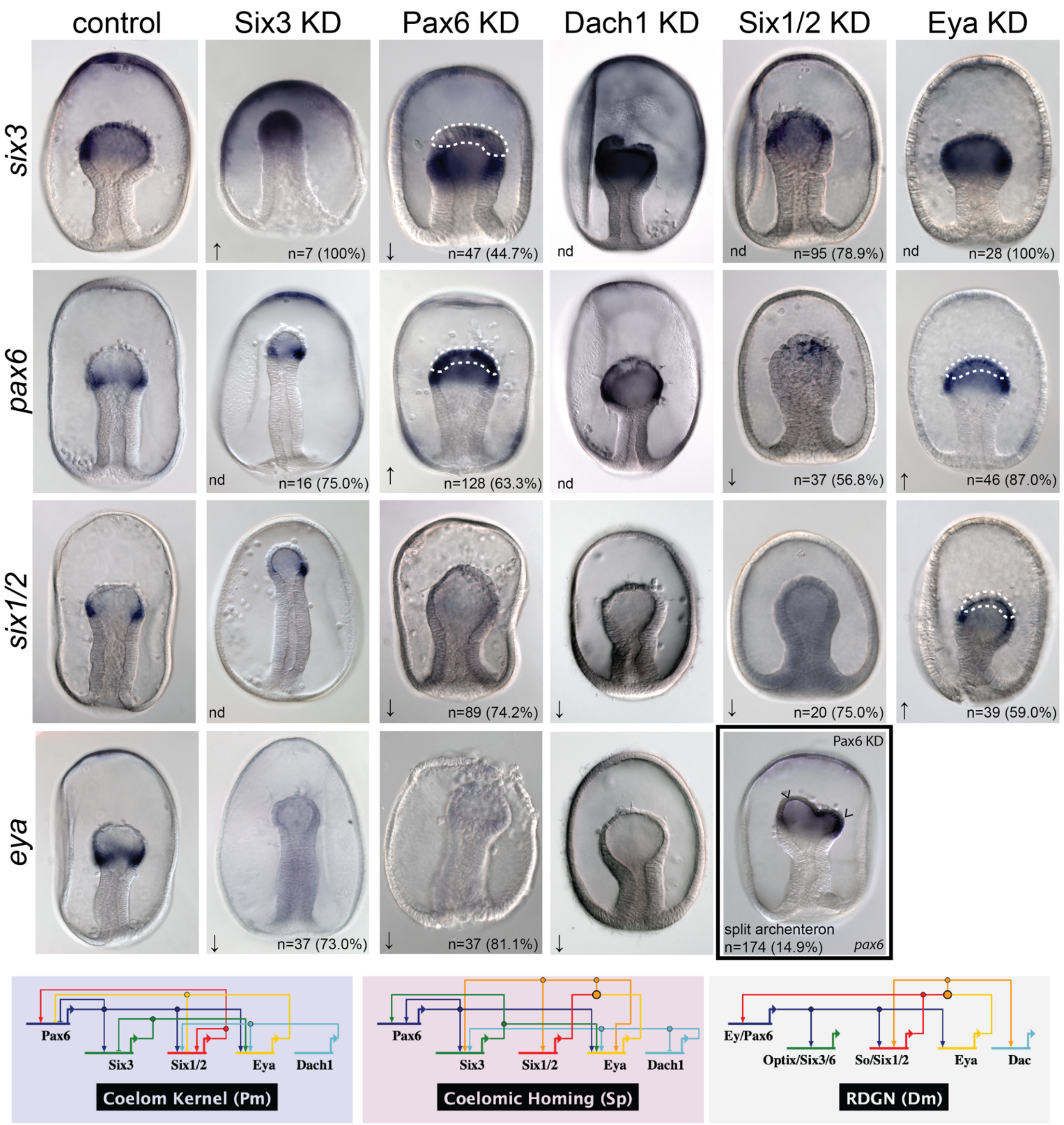
control
Six3 KD
Pax6 KD
Dach1 KD
Six1/2 KD
Eya KD
six3
n=7 (100%)
n=47 (44.7%)
nd
nd
n=95 (78.9%)
nd
n=28 (100%)
pax6
nd
n=16 (75.0%)
n=128 (63.3%)
nd
n=37 (56.8%)
n=46 (87.0%)
six1/2
nd
n=89 (74.2%)
n=20 (75.0%)
n=39 (59.0%)
eya
n=37 (73.0%)
n=37 (81.1%)
Pax6 KD
split archenteron
n=174 (14.9%)
pax6
Pax6
Six3
Six1/2
Eya
Dach1
Coelom Kernel (Pm)
Pax6
Six3
Six1/2
Eya
Dach1
Coelomic Homing (Sp)
Ey/Pax6
Optix/Six3/6
So/Six1/2
Eya
Dac
RDGN (Dm)

**Figure 8.** Subcircuit including *pax6*, *six3*, *eya*, *dach1*, and *six1/2* involved in coelomogenesis in echinoderms and is similar to Drosophila retinal determination network. Data shown are all colorimetric WMISH using the probes indicated on the left in the conditions listed along the top. *six3* expression is normally distributed throughout the mesodermal bulb of the archenteron at 48 hpf, while *pax6*, *six1/2*, *dach1*, and *eya* are normally expressed at the posterior aspect of the mesodermal bulb, having been cleared earlier from anterior regions of the mesoderm. Phenotypic effect of the perturbation of each gene is indicated, including no difference (nd), increase (↑), decrease (↓). The number of embryos assessed and percent of embryos expressing the phenotype are also reported. Some reported phenotypes are localized to the top of the archenteron and are highlighted (dashed line); e.g. the effect of Pax6 knockdown on *six3* expression is reported specifically for the anterior region of archenteron (dashed line). Some Pax6 knockdown embryos exhibited a bifurcated archenteron (e.g. boxed panel, “split archenteron”). These results enabled us to construct a network sea star coelom and are connected in a similar regulatory sub-network in both sea star and sea urchin coelomic mesoderm, which is strikingly similar to the retinal determination gene network (RDGN) in Drosophila.

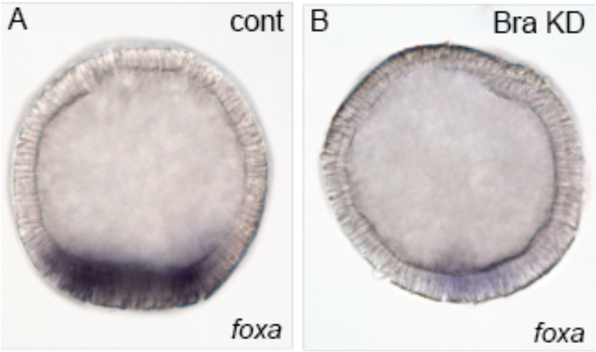


**Figure 9.** Knockdown of *foxA* transcript by morpholino oligonucleotide targeting the Bra transcript (B) compared to control morpholino injection (A) by colorimetric WMISH.

**Table S1.** Morpholino antisense oligonucleotide sequences and concentrations used.

| Morpholino | Sequence | Injection Solution Concentration |
|---|---|---|
| Tgif_1 | TCGAGAGCCGATCCACGAATCAAGA | 800 µM |
| Tgif_2 | TGCAACAACAAAGACGGTTCGGTAC | 400 µM |
| HesC_1 | CAATCATGCTGAAGATTGTCGAAGG | 600 µM |
| HesC_2 | TGTTCCGAGTAGAAGACGAATCGA | 600 µM |
| Six3_1 | ACATTGAGCCGAGCATCTGGACCCG | 600 µM |
| Six3_2 | TCTCAGCAGCGCAGTCGAGAGACAC | 600 µM |
| Pax6_1 | AAGTGCTTCACTGACCTGTATCCTA | 400 µM |
| Pax6_2 | CTCTGAAGTAAACTGTGTATAAGGC | 600 µM |
| Eya_1 | GCACGTAGTTGAAGCAAACACATCA | 600 µM |
| Six1/2_1 | CGTGAAGCCAAACGACGGCAACATG | 600 µM |
| Krox_1 | CAGGTCCTTTCATTCTGGTACTCAG | 600 µM |
| Bra_1 | CACTCATGGTGTTTCAAAAATGCTC | 600 µM |
| Standard Control Oligo | CCTCTTACCTCAGTTACAATTTATA | 600 µM |

**Table S2.** Primers used for quantitative RT-PCR assessment of knockdown and drug treatment conditions.

| Gene | F | R |
|---|---|---|
| *Hesc* | AAGCCTCATCTTCCCAGCTCTC | CCTTCAGGTAACGGACGGTCAT |
| *Delta* | CGAAGGCTTCACGTGCTACTG | GCGCATGCGTAGCCATTCTC |
| *lamin2b receptor* | GAGCATGCCTAAGCCAGACC | CTCCACCATGGGCTCCAGTA |

**Table S3.** Descriptions and phenotype counts for morpholinos to coelomic network genes.

| Morpholino | WMISH Probe | Phenotype | # Counted | # Phenotypic | % phenotypic |
|---|---|---|---|---|---|
| Pax6 | Eya | Decrease | 37 | 30 | 81.1% |
| Six3 | Eya | Decrease | 37 | 27 | 73.0% |
| Pax6 | GataE | no mesodermal expression | 48 | 4 | 8.3% |
| Eya | GataE | no mesodermal expression | 45 | 34 | 75.6% |
| Six1/2 | GataE | no mesodermal expression | 44 | 20 | 45.5% |
| Eya | Pax6 | clearance loss | 46 | 40 | 87.0% |
| Pax6 | Pax6 | clearance loss | 128 | 81 | 63.3% |
| Six1/2 | Pax6 | Decrease | 37 | 21 | 56.8% |
| Pax6 | Six1/2 | Decrease | 89 | 66 | 74.2% |
| Eya | Six3 | normal expr | 28 | 28 | 100.0% |
| Pax6 | Six3 | Off in anterior mesoderm | 47 | 21 | 44.7% |
| Six1/2 | Six3 | normal expr | 95 | 75 | 78.9% |
| Six3 | Six3 | Increase | 7 | 7 | 100.0% |
| Pax6 | -ALL- | split archenteron | 174 | 26 | 14.9% |
